# Neural Mechanisms of Cognitive Flexibility and Interference Control in a Stroop Task Switching Paradigm

**DOI:** 10.1101/2025.05.05.652181

**Authors:** Zeynab Tahamtan, Seyed Ali Osia, Pawel Herman

## Abstract

Task-switching paradigms, often used to study cognitive flexibility, frequently employ incongruent bivalent stimuli, triggering two tasks and potentially conflating cognitive flexibility with interference control. This study assesses cognitive flexibility using univalent stimuli (triggering one task) and congruent bivalent stimuli (same response across tasks) in a modified Stroop task to investigate well-established neural activity correlates of cognitive flexibility, manifested in human (females and males) electroencephalography (EEG) recordings, while isolating switch process from the influence of interference control at the response level. In particular, we analyzed EEG theta-band activity and event-related potential (ERP) components in switch (N2, P3a, P3b, late sustained potential (LSP)) and interference (N400, LSP and mid-frontal theta activity) conditions. We compared each of the switch and the interference condition to the control condition using cluster-based permutation test. In the switch condition, we observed fronto-central N2, reduced frontal P3a, and a positive occipital LSP. The interference condition showed increased frontal theta, parietal N400, and a positive occipital LSP. We also compared switch and interference conditions using cluster-based permutation test. We observed a larger N2 in the frontocentral regions during the switch condition and higher frontal theta activity during the interference condition, which aligns with their comparisons to the control condition. This result suggests that distinct neural mechanisms are used for each of the processes involved in conflict monitoring. Specifically, the theta activity may reflect sustained monitoring and conflict resolution during interference, while the N2 may reflect more transient conflict detection and the need to switch task sets.

**Significance Statement:** Cognitive flexibility—the ability to adapt behavior to a changing environment—is essential for goal-directed actions. For assessing cognitive flexibility unlike the typical approach, we did not use incongruent bivalent stimuli. We used univalent and congruent bivalent stimuli to isolate cognitive flexibility while assessing interference control separately with incongruent bivalent stimuli. We analyzed well-documented brain activities related to cognitive flexibility (P3b, N2, P3a, LSP) and interference control (N400, LSP), and mid-frontal EEG theta activity. Despite using different stimuli, we observed all expected components associated with the switch process except for the P3b. Both processes share common parietal activity, and while the frontal lobe plays a role in both, its activity differs between them.

## Introduction

Cognitive flexibility is the ability to adapt behaviors to a changing environment and switch between different tasks (Scott, 1962). It is a core component of executive functions, along with other top-down functions such as inhibition and working memory (Diamond, 2013). These functions are all involved in goal-directed behavior, as opposed to habitual responses, but each function has a specific component that separates it from the others, suggesting both unity and diversity among executive functions (Teuber, 1972; Friedman et al., 2008). The unique component in cognitive flexibility is “shifting”, which distinguishes it from general executive function capabilities (Miyake & Friedman, 2013). Although these functions are distinct, experimental paradigms typically used to evaluate cognitive flexibility often involve interference control, which is a subcomponent of inhibition (Diamond, 2013). These experiments build on task-switching paradigms, in which individuals alternate between tasks (Dajani & Uddin, 2015).

Shifting between tasks elicits a “switch cost” effect, wherein participants respond more slowly and commit more errors when alternating between tasks compared to repeating the same task (Jersild, 1927). Nearly all experiments that use task-switching paradigms involve incongruent bivalent stimuli (Diamond, 2013). These stimuli encode at least two rules, each linked to a specific stimulus dimension, where the correct response for one rule is incorrect for the other (Grundy et al, 2013). Consequently, when the task changes, participants are forced not only to shift between tasks but also to suppress attention to one stimulus dimension, in order to manage interference. Hence, assessing cognitive flexibility independently from interference control is challenging.

To overcome this challenge, this study employed a task-switching paradigm using univalent stimuli and congruent bivalent stimuli. This approach eliminates response conflict as only one response rule can be applied to these stimuli. We did not use cues given that cues might introduce bias (Logan & Bundesen, 2003; Mayr & Kliegl, 2003) and cues are optional for univalent stimuli and do not reduce switch costs (Elchlepp et al., 2012). When assessing cognitive flexibility we exclude incongruent bivalent stimuli, in contrast to many Stroop task-switching studies that have solely focused on them (e.g, Wylie & Allport, 2000; Monsell et al., 2000). This approach is expected to minimize interference control and emphasize the shifting process, a key indicator of cognitive flexibility, as the dominant process in switch trials.

In our modified Stroop task, participants (9 females and 11 males) performed three conditions based on stimuli and rules designed to investigate distinct cognitive processes: switch, interference, and control. The switch condition involved participants alternating between rules without the influence of interference control at the response level. The interference condition aimed to assess the interference effect without the influence of task-switching. The control condition served as a baseline where neither interference nor switching was present, allowing us to compare the neural activity in the other conditions to a neutral state.

To quantify interference and switch effects, we measured RT, ACC and recorded electroencephalography (EEG) from the participants performing the task and examined well-established event-related potentials (ERP) components associated with task-switching (P3b, N2, P3a, LSP) (Gajewski & Falkenstein, 2011; Karayanidis & Jamadar, 2014; Berti, 2016; Brass et al., 2005) and interference control (N400, LSP) as well as mid-frontal EEG theta activity (Appelbaum et al., 2009; Badzakova-Trajkov et al., 2009, West., 2003, Tang et al., 2013; Hanslmayr et al., 2008). We anticipated both shared and distinct neural patterns in the switch and interference conditions, given their roles in executive functions. We expected to observe well-established ERP components associated with task-switching in the switch condition despite excluding incongruent bivalent stimuli which are typically employed in task-switching. Frontal lobe activity differed by task (switch: anterior N2; interference: increased theta-band activity), while parietal LSP was shared. Compared to control, the switch condition showed all expected cognitive flexibility components except P3b.

## Materials and Methods

### Participants

Twenty volunteers (11 females, 9 males; average age = 28.05 years, SD = 3.30) from the local community participated. All participants had normal or corrected-to-normal vision, and no history of neurological disorders. Informed consent was obtained from all participants, they were instructed to get sufficient sleep and avoid alcohol for 24 hours before the study. The data from one of these subjects was subsequently omitted from analysis because of low quality of the EEG signal.

### Experimental Design

Participants completed a modified version of Stroop color-word test (Stroop,1935), which was adapted and modified from a previous study (West & Alain, 1999). Participants were presented with color words (“red,” “green,” “blue,” and “yellow”) and instructed to select the right color among four possible options. They should respond by pressing one of the four keys (i.e., V, B, N, or M) on a computer keyboard, corresponding to the different colors, using the index and middle fingers of both hands. They were required to respond as quickly and accurately as possible.

The classic Stroop test consists of three kinds of stimuli; congruent, incongruent, and neutral. In the congruent stimuli, the color words matched the ink color (e.g., the word “yellow” displayed in yellow ink). In the incongruent stimuli, the color words were presented in a different ink color (e.g., the word “yellow” displayed in green ink). In the neutral stimuli, non-color words were displayed in red, blue, green or yellow ink colors (e.g., “bird” displayed in yellow ink). For these stimuli, participants were instructed to respond based on the ink (color-based rule) e.g. when the ink color is red, the participant should select red, regardless of the written word, whether its “blue”, “red” or “bird”. In addition to these usual stimuli in the classic Stroop test, we introduced a fourth category of stimuli, gray stimuli, where the color words were presented in gray ink (e.g., the word “yellow” displayed in gray ink). For these stimuli, participants were required to change their response strategy and focus on the semantic meaning of the word, and respond based on the word (word-based rule). The experiment was initiated after the rules were presented. Once participants had read the instruction, they pressed the spacebar to indicate their readiness to proceed. The stimuli were presented at the center of the screen for 400 ms, followed by a fixation cross (“+” symbol) that remained until a response was made. After the participant’s response, there was an intertrial interval of 1000 ms.

The experiment was conducted in three phases. The initial phase, referred to as the “color-to-key acquisition phase,” consisted exclusively of congruent stimuli and included 100 trials, with each color word presented 25 times in random order. This phase aimed to establish a strong mapping between the stimulus colors and response keys, ensuring that participants effectively learned the key associated with each color. The second phase, known as the “Practice Phase,” comprised 40 trials representing four types of stimuli: congruent, incongruent, neutral, and gray. Each stimulus type was presented 10 times in random order. This phase aimed to provide participants with practice in responding to various stimuli. After each response, feedback was provided for 1000 ms, indicating whether the response was correct or incorrect. Feedback was given only during the practice phase.

The third phase, referred to as the “Test Phase,” included all trial types (congruent, neutral, incongruent, and gray). These stimuli were presented in three blocks, each varying in the proportion of gray stimuli, with the random order of block presentation and a self-paced break between each block. The “zero gray” block consisted of 96 trials, exclusively presenting congruent and incongruent stimuli. The “one forth gray” block also included 96 trials, with each type of stimulus accounting for 25% of all trials. The “half gray” block consisted of 144 trials, with an additional 48 trials assigned to gray stimuli, resulting in 50% of the trials being gray stimuli. Our analysis focused exclusively on the Test Phase.

To assess cognitive flexibility and interference control during the Test Phase, we defined three conditions. In the Switch Condition, we focus on univalent stimuli and congruent bivalent stimuli in switch trials, where participants alternate between rules. This approach eliminates interference by excluding incongruent bivalent stimuli from the analysis. The Interference Condition involves the use of only incongruent stimuli in repeat trials, where the same rule is applied as in the previous trial. This condition specifically targets interference control without the influence of task switching. Finally, the Control Condition consists of repeat trials featuring congruent bivalent and neutral stimuli. This ensures that there is no impact from either interference or switching. An overview of the stimuli and trial types, including the color-based and word-based rules, is illustrated in Figure 1. In the “zero gray” block, we wanted to ensure that we had enough incongruent trials, which were also repeated trials, since there were no gray trials and no switches in this block. In the “half gray” block, we aimed to ensure that there were enough switching trials, as there was a larger proportion of gray trials, requiring participants to adapt to a new rule. Also, the “one-fourth gray” block consists of more control trials while maintaining switch and interference trials.

**Figure 1.**
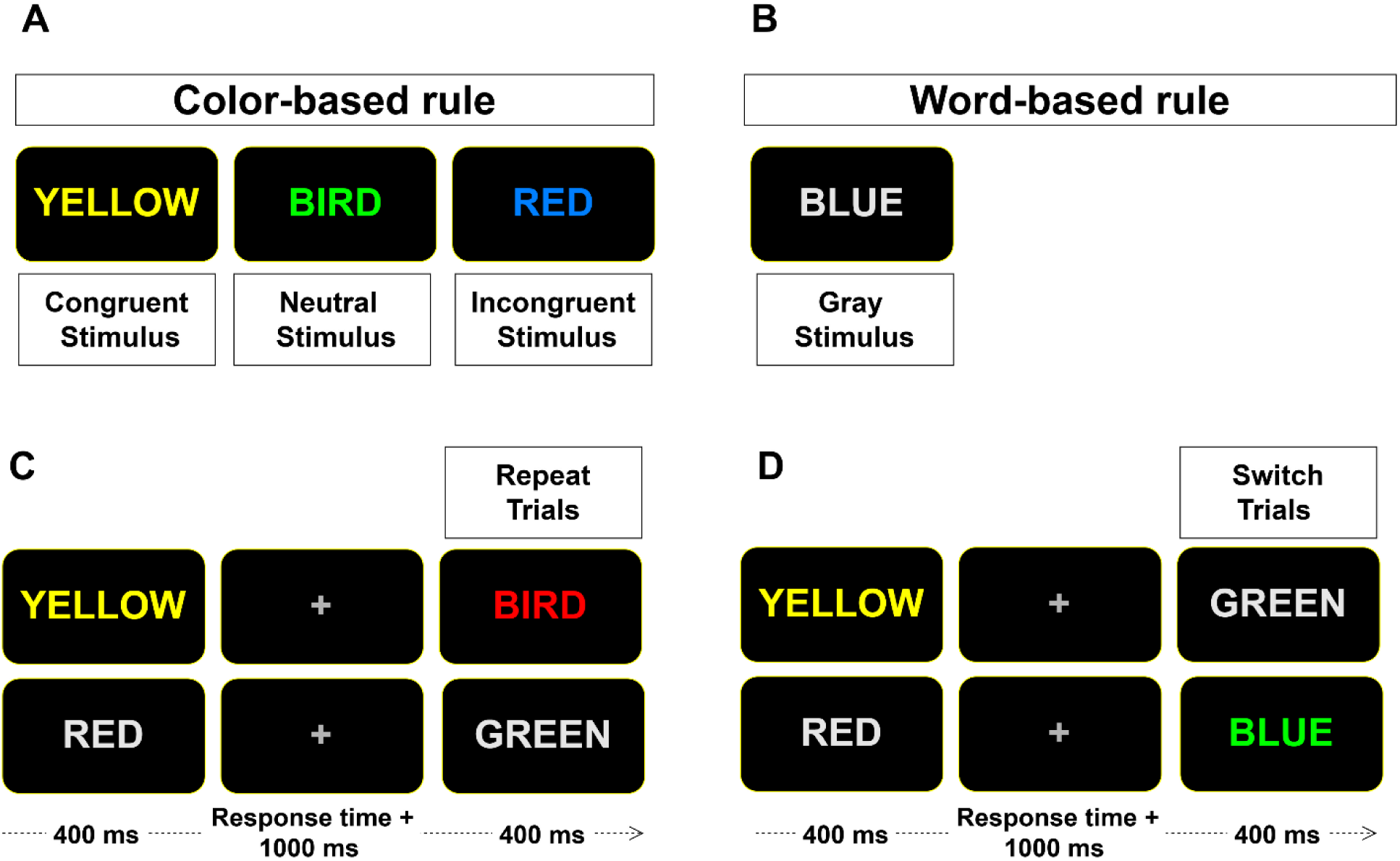
Overview of Experimental Design and Trial Types. The experiment involved two rule types: a color-based rule and a word-based rule. In the color-based rule, participants responded to the ink color of the stimuli (A). Three trial types were used: congruent stimuli, neutral stimuli, and incongruent stimuli. In the word-based rule, participants responded to the semantic meaning of the word, with gray stimuli(B). The experimental design includes repeat trials, where the rule remained consistent with the previous trial (e.g., consecutive color-based trials) (C). Switch trials required participants to switch from the rule used in the previous trial to a different rule (e.g., switching from a word-based to a color-based rule) (D). Each stimulus was presented for 400 ms, followed by a fixation cross that remained until a response was made. A 1000 ms interval followed each response before the next trial began.

The experiment was programmed with MATLAB software (MathWorks, Inc., Natick, MA, USA) and the Psychophysics Toolbox Version 3 (Psychtoolbox-3; Brainard, 1997; Pelli, 1997; Kleiner et al., 2007).

### Recording and Preprocessing

The EEG was recorded from Cognionics (CGX) Quick-20r v2 headset, a 21-channel, dry, wireless EEG device with sampling rate of 500 Hz at standard 10/20 locations (Fp1, Fp2, F7, F3, Fz, F4, F8, T7, C3, Cz, C4, T8, P7, P3, Pz, P4, P8, O1, O2). The electrode impedance was maintained below 300 kΩ. The signals were recorded using a left earlobe reference electrode and then re-referenced offline to the mean of the left and right earlobes and filtered with a bandpass of 0.5 – 40 Hz.

Independent component analysis (ICA) was subsequently conducted for each participant to identify and remove components explicitly linked to eye blinks. For each participant, the cleanest data segment from the test phase was selected for ICA decomposition using the FastICA algorithm (Ablin et al., 2018). A trained expert then inspected the components and removed those associated with blink artifacts across the entire test phase dataset. Following artifact correction, the EEG data were epoched from 1000 ms before stimulus onset to 1400 ms after stimulus onset for further analysis. We focused exclusively on correct trials from the test phase, defined as trials in which participants provided the correct response within a response time of less than 1500 ms.

After epoching, several techniques were implemented to exclude signals containing significant muscle artifacts, extreme voltage offsets, flat signals, and bad channels. If a channel exhibited any of these issues, we labeled them as bad channels in each epoch and excluded them from the rest of the analyses without excluding the entire epoch. To identify extreme voltage offsets and flat signals, channels with a peak-to-peak amplitude exceeding 90 μV or below 1 μV were flagged and removed from further analysis to ensure data quality during preprocessing. Brain signals typically exhibit skewed (often lognormal) distribution patterns and demonstrate a nearly linear decline in power as frequency increases on a logarithmic scale (Buzsáki & Mizuseki, 2014). In contrast, bad channels and signals with a high noise ratio display distinct distribution patterns or maintain a constant spectrum lacking a prominent frequency, often referred to as ’white noise’ (Barry & Blasio, 2021; Keil et al., 2022). This type of noise may arise from random external influences on the brain or intrinsic noise during signal recording and digitization, potentially including inaccuracies during analog-to-digital conversion (Keil et al., 2022). To eliminate noisy channels with significant muscle artifacts and high noise ratios, which exhibit distribution patterns differing from brain signals, a linear regression approach was employed to model the logarithmically transformed power spectral density (PSD) data (Fitzgibbon et al., 2019). Using this approach, we identified channels that do not follow the lognormal pattern and label them as noisy.

After applying various techniques to detect and label noisy channels, we proceeded with further analysis on the remaining 80% of signals, which were used for ERP and time-frequency analyses. During epoch averaging, channels identified as noisy were excluded from the analysis; only noise-free channels were considered to ensure the integrity of the data.

### Statistical Analysis

#### Behavioral

For Behavioral analysis we compared reaction time (RT) and response accuracy (ACC) among three conditions (switch, interference, and control). Response accuracy was calculated as the total number of correct responses divided by the total number of responses (correct and incorrect) for each participant. For RT analysis, only trials with correct responses were included. Mean reaction times and response accuracies were calculated for each participant and condition.

Repeated measures ANOVA were conducted to compare RTs and ACCs across the three conditions. For post-hoc analysis, pairwise t-tests were performed with false discovery rate correction using the Benjamini-Hochberg (FDR-BH) method. To assess the switch cost, RT and ACC in the switch condition were compared to those in the control condition. Similarly, to evaluate the interference effect, RT and ACC in the interference condition were compared to those in the control condition. Additionally, we compare RT and ACC in the switch condition with interference control.

#### EEG correlates: ERPs and oscillations

For the ERP analysis, we used a time range of -200 ms to 800 ms relative to the stimulus onset. Baseline correction was performed for each trial using the pre-stimulus interval (−200 ms to 0 ms). To examine differences between conditions for specific candidate components (e.g. N2), we selected relevant channels and time windows based on the existing literature. Spatio-temporal cluster-based permutation tests (Maris & Oostenveld, 2007) were used to evaluate significant differences between conditions, addressing the multiple comparisons problem. For each comparison, univariate t-tests were performed at every sample in time and then the set of neighboring points associated with a first-level significant p-value are considered as candidate clusters (Candia-Rivera & Valenza 2022). For this phase, we used a liberal p-value threshold of 0.1 to capture weak and widespread effects, ensuring larger clusters without compromising the validity of the cluster-based permutation test (Maris & Oostenveld, 2007). Candidate clusters are then quantified based on their mass i.e. sum of t-values and then significant clusters with p-values smaller than 0.05 are identified based on a permutation test on mass values. We have used 2,000 randomizations as it is common to use Monte Carlo estimation of the permutation distribution and the number of randomizations is recommended to be more than 1,000 (Meyer et. al 2021). For defining spatio-temporal neighborhood in each brain region, electrodes within about ∼5 cm of one another were considered spatial neighbors (Meyer et al. 2021) and adjacent time points were considered temporal neighbors. Finally, we computed the grand average ERP by averaging across participants for each condition and channel, which was used for all subsequent ERP plots.

For the time-frequency analysis, we focused exclusively on the theta activity on the frontal region (F3, Fz, F4). We used a time range of -450 ms to 800 ms relative to stimulus onset. For each participant and condition, time-frequency representations (TFRs) were calculated for all clean trials individually with the support of multitaper spectral analysis. We employed Discrete Prolate Spheroidal Sequences (DPSS) tapers generated with the Slepian sequence. The frequency range of interest was 3.5-7.5 Hz with a frequency resolution of 1 Hz and half of cycles per frequency was used to balance temporal and frequency resolution. The TFRs were averaged across clean trials for each participant and condition, followed by baseline correction. Baseline correction based on pre-stimulus baseline (−450 ms to -50 ms) was performed by calculating the logarithm of ratio of the power at each time point to the average power in the pre-stimulus baseline interval, and results were expressed in decibels (dB). We averaged the TFRs from the F3, Fz, and F4 channels at the end and conducted cluster-based permutation tests to detect significant differences between conditions. The parameters for the cluster-based permutation test were similar to those used in the ERP test mentioned above.

To compare all three conditions (switch, interference, and control), cluster-based permutation tests were performed on well-established ERP components and TFRs associated with cognitive flexibility (N2, P3a, P3b, LSP) and interference control (N400, LSP, and frontal theta activity). These analyses were conducted at specific time intervals and brain regions based on prior knowledge. For the N2 and P3a components, tests were conducted within a 200–500 ms time window at frontal (F3, Fz, F4) and central (C3, Cz, C4) electrode sites (Gajewski & Falkenstein, 2011; Wu, 2015). For the P3b component, tests were performed at parietal sites (P3, Pz, P4, P7, P8) within a 300–500 ms time window (Gajewski & Falkenstein, 2011; Wu, 2015). For the LSP, tests were conducted at parietal and occipital (O1, O2) regions within a 500–800 ms time window, consistent with previous studies on cognitive flexibility (Brass et al., 2005; Wu, 2015) and interference control (Appelbaum et al., 2009; Donohue et al., 2016; Markela-Lerenc et al., 2009; West, 2003). The N400 component was tested at parietal sites within a 400–500 ms time window (Donohue et al., 2016; Hanslmayr et al., 2008; Larson et al., 2009; Liotti et al., 2000; Markela-Lerenc et al., 2004; Qiu et al., 2006; Rebai et al., 1997). Finally, frontal theta activity was examined, within 400-800 ms in averaged frontal TFRs (Atchley et al., 2017; Cohen, 2014; Ergen et al., 2014; Hanslmayr et al., 2008; Tang et al., 2013).

## RESULTS

We investigated cognitive flexibility in a switch condition by excluding incongruent stimuli in switch trials to eliminate response-level interference. Additionally, we examined interference control in a separate interference condition, traditionally assessed using incongruent stimuli. A control group, without switch or interference, was included for comparison with both experimental conditions. To investigate the cognitive processes underlying interference control and cognitive flexibility, we analyzed behavioral data measuring RT and ACC, and EEG correlates to quantify effects of interference and switch.

### Behavioral

In our study we measured RT as the time elapsed between the presentation of the stimulus and the participant’s response. For RT analysis, only trials with correct responses were included. ACC was quantified as the percentage of correct responses. To compare these behavioral measures across the conditions, we applied repeated measures ANOVA.

For RT (Fig. 2), the analysis revealed a significant main effect of condition, (F(2, 36) = 26.52, p = 7e-6). Post-hoc pairwise t-tests with FDR-BH correction showed that RTs were significantly slower in the interference condition compared to control (t(18) = -7.02, p-corrected < 1e-5), and in the switch condition compared to control (t(18) = -3.86, p-corrected = 0.001). RTs were also significantly slower in the interference condition compared to switch (t(18) = 3.80, p-corrected = 0.001).

**Figure 2.**
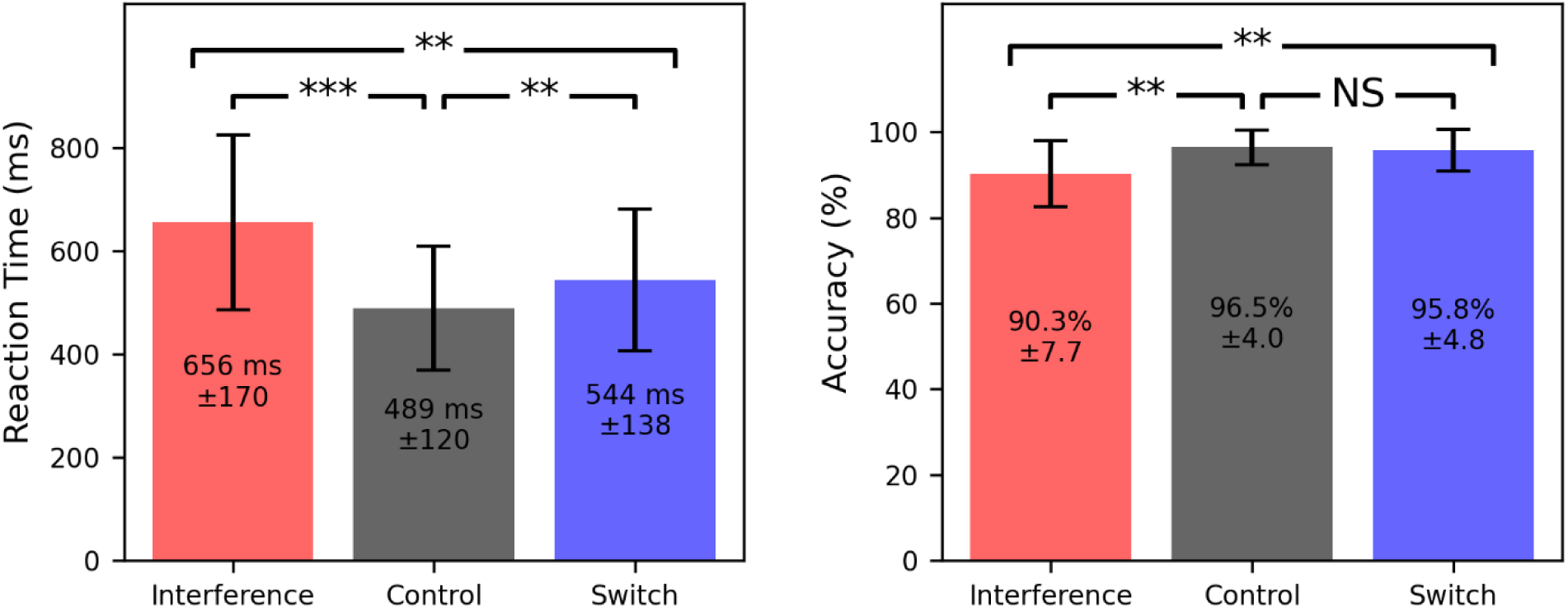
RT (left panel) and ACC (right panel) across three experimental conditions. The charts reflect mean RTs and ACCs across the three conditions: interference, control and switch conditions. The mean values are accompanied by the error bars reflecting the standard error across the experimental conditions. Significant differences were identified using Repeated measures ANOVA. RTs were significantly slower in the Switch and Interference conditions compared to the Control condition and the Switch compared to the Interference conditions. ACC was significantly lower in the Interference condition compared to the Control (*p < 0.05, ** p < 0.01, *** p < 0.001).

For ACC, a significant main effect of condition was also found (F(2, 36) = 11.99, p = 1.02e-4). Post-hoc pairwise comparisons revealed that accuracy was significantly lower in the interference condition compared to control, (t(18) = 3.57, p-corrected = 0.003), and in the interference condition compared to switch, (t(18) = -3.55, p-corrected = 0.003). The comparison between control and switch was not statistically significant, (t(18) = 1.21, p-corrected = 0.24).

### EEG correlates: ERPs and oscillations

We conducted cluster-based permutation tests within all three conditions (switch, interference, and control) to assess discriminative aspects of well-established ERP components and oscillatory correlates of cognitive flexibility (N2, P3a, P3b, LSP) and interference control (N400, LSP, and frontal theta activity). Although we focused our analysis on a predefined time window and region of interest, we do not report the specific boundaries of the significant cluster identified through our cluster-based permutation test because cluster-based permutation tests do not provide precise information about the exact temporal or spatial extent of the effect (Sassenhagen & Draschkow, 2019).

Testing for an N2 and P3a effect in the latency range from 200 to 500 ms post-stimulus in frontal region, the cluster-based permutation test revealed a significantly more negative amplitude in the switch condition compared to both the control (p = 5e-4, 19 subjects, 2000 permutations) and interference (p = 0.02, 19 subjects, 2000 permutations) conditions. In this latency range, for N2 and P3a the difference was located in frontal and most pronounced over left frontal sensors.

There was no significant difference between control and interference conditions (p > 0.05, 19 subjects, 2000 permutations) (Fig. 3A). Figure 3B displays the grand average ERP with maximum effect in the negative cluster in the frontal site (F3), where both the N2 and P3a components are present.

**Figure 3.**
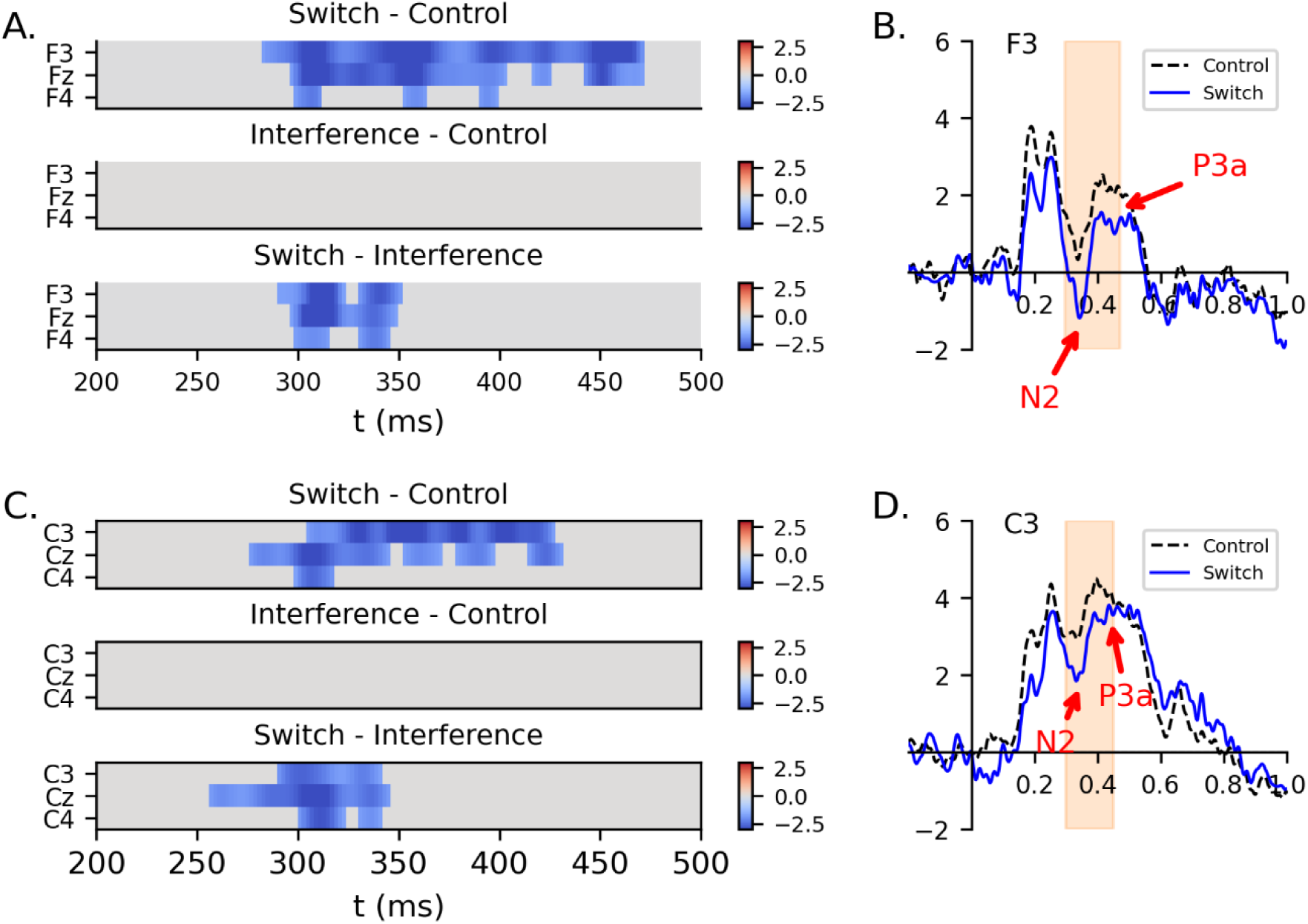
N2 and P3a component in the frontal and central region. The raster diagram shows a negative cluster between ERPs for the switch vs. control and switch vs. interference conditions, within the 200-500 ms time window in the frontal (A) and central (C) site. Time is expressed relative to the stimulus onset. Grand average ERP waveforms for electrodes F3 (B) and C3 (D) illustrate the channels with the strongest effects in these negative clusters. The orange shaded areas reflect the time window falling into the negative cluster. There is no significant difference for interference vs. control comparison.

For N2, a similar pattern was observed in the central region (Fig. 3C) with a significant difference between switch and control (p = 0.001, 19 subjects, 2000 permutations), and switch and interference (p = 0.008, 19 subjects, 2000 permutations) conditions. Fig. 3D displays the grand average ERP at electrode C3, which showed the strongest effect within the negative cluster at central sites. There was no significant difference between control and interference conditions (p > 0.05, 19 subjects, 2000 permutations) (Fig. 3C).

Testing for an N400 effect in the latency range from 400 to 500 ms post-stimulus in the parietal site, the cluster-based permutation test revealed significantly more negative amplitude between the interference and the control condition (p = 0.04, 19 subjects, 2000 permutations) shown in

Fig. 4A. However, no significant differences were found between the switch condition compared to control and interference conditions (p > 0.05, 19 subjects, 2000 permutations). Fig. 4B displays the grand average ERP at electrode P4, showing the strongest effect within the negative cluster.

**Figure 4.**
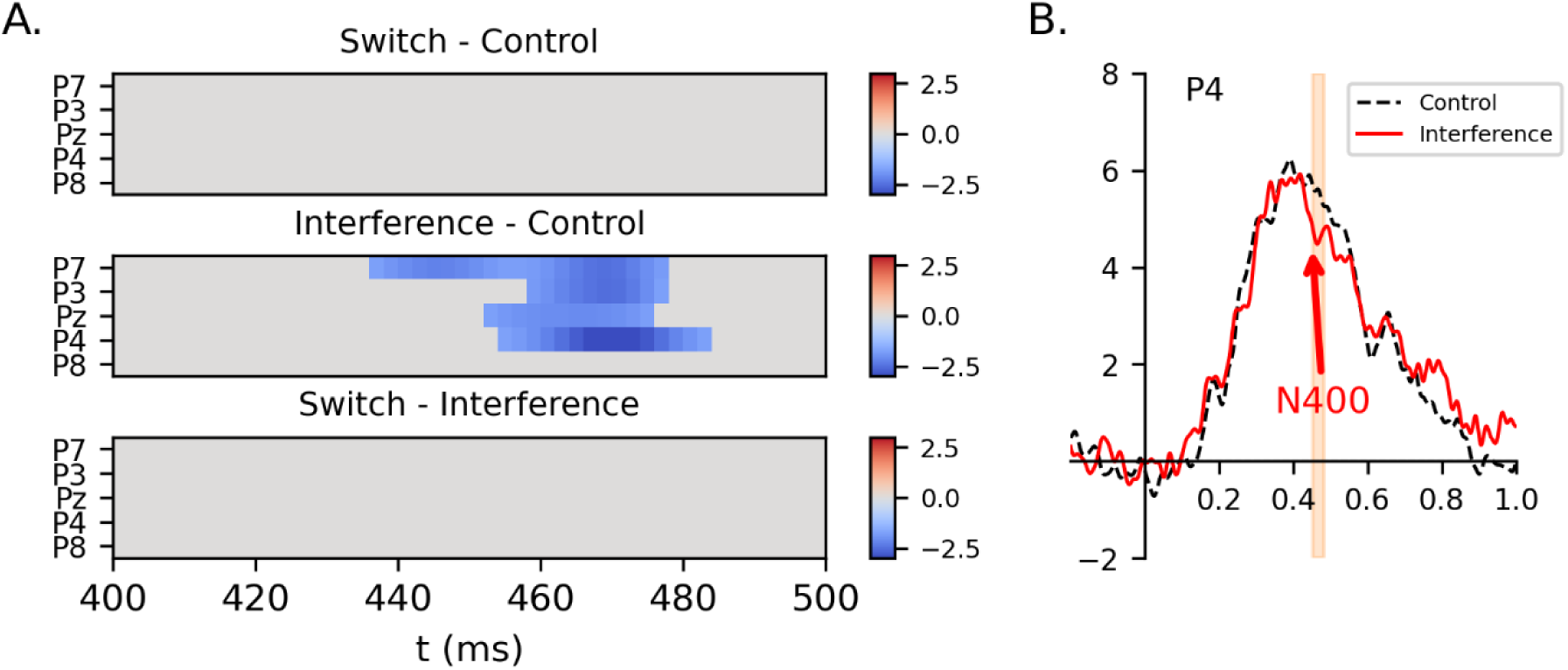
N400 component in the parietal region. The raster diagram shows a negative cluster between ERPs for the interference vs control conditions, within the 400-500 ms time window in the parietal site (A). Time is expressed relative to the stimulus onset. Grand average ERP waveform for electrode P4 illustrates the channel with the strongest effects in these negative clusters (B). The x-axis depicts time (seconds) relative to stimulus onset. The orange shaded areas reflect the time window falling into the negative cluster. There is no significant difference for switch vs. control and switch vs. interference comparisons.

Testing for a positive LSP effect in the latency range from 500 to 800 ms post-stimulus in the parietal site and occipital site, the cluster-based permutation test revealed significantly more positive amplitude difference between the switch (p = 5e-4, 19 subjects, 2000 permutations) and interference (p = 5e-4, 19 subjects, 2000 permutations) conditions and the control condition in occipital site. There were no significant differences between switch and interference conditions (p > 0.05, 19 subjects, 2000 permutations) (Fig. 5A). Additionally, Fig. 5B displays the grand average ERP at electrode O2, showing the strongest effect within the positive cluster at occipital sites.

**Figure 5.**
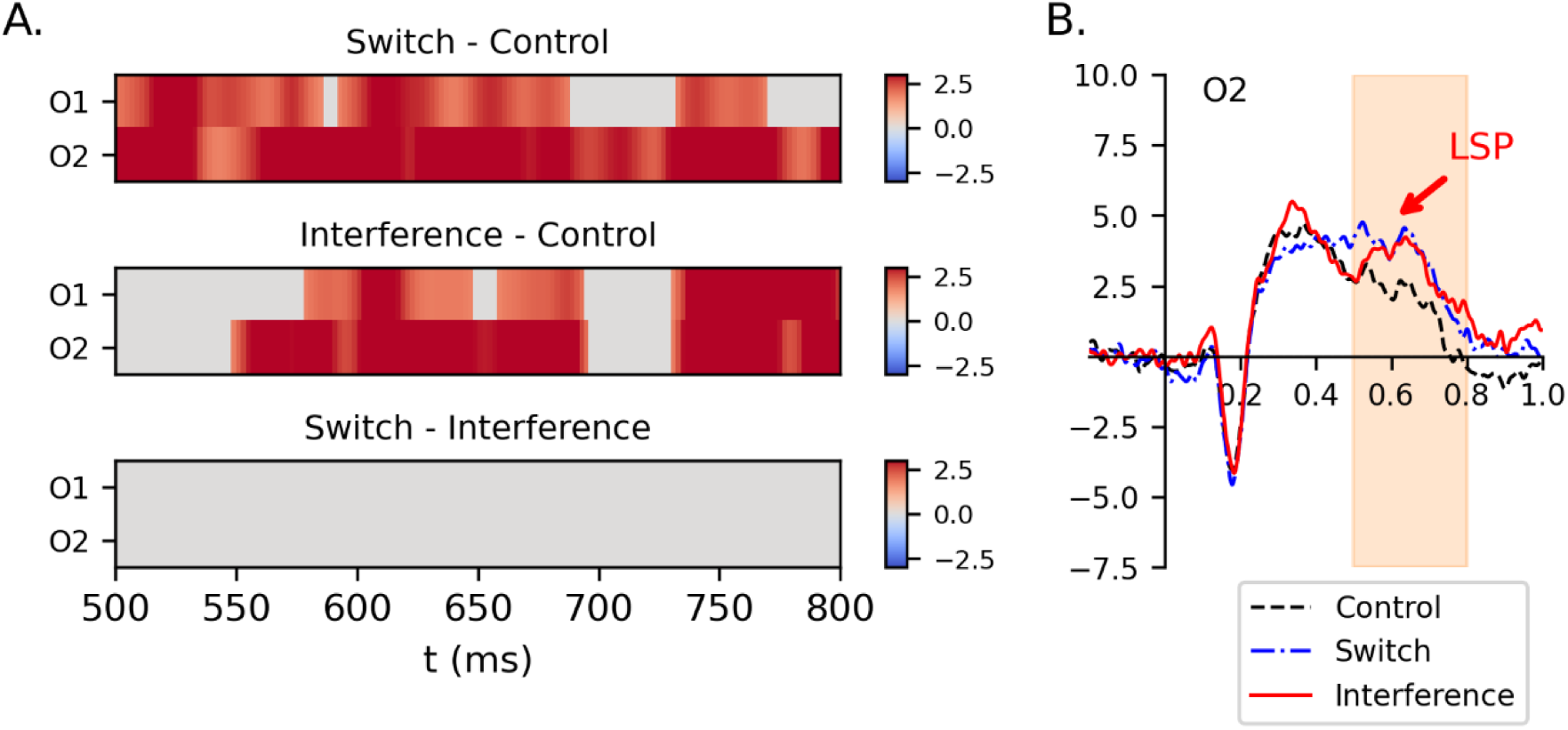
LSP component in the occipital region. The raster diagram shows a positive cluster between ERPs for the interference vs. control and switch vs. control conditions within the 500-800 ms time window in the occipital site (A). Time is expressed relative to the stimulus onset. Grand average ERP waveform for electrode O2 illustrates the channel with the strongest effects in this positive cluster (B). The x-axis depicts time (seconds) relative to stimulus onset. The orange shaded areas reflect the time window falling into the positive cluster. There is no significant difference between interference vs. switch comparison.

To examine well-established brain activity associated with interference control, we also analyzed discriminative correlates in the frontal theta band oscillations. To this end, we conducted a cluster-based permutation test on theta band power (3.5 - 7.5 Hz) in the averaged TFRs of the frontal site. We observed differences between the interference and the control condition (p = 0.04, 19 subjects, 2000 permutations) and between the interference and the switch condition (p = 0.01, 19 subjects, 2000 permutations). However, there were no significant differences between the switch and control condition (p > 0.05, 19 subjects, 2000 permutations) (Fig. 6).

**Figure 6.**
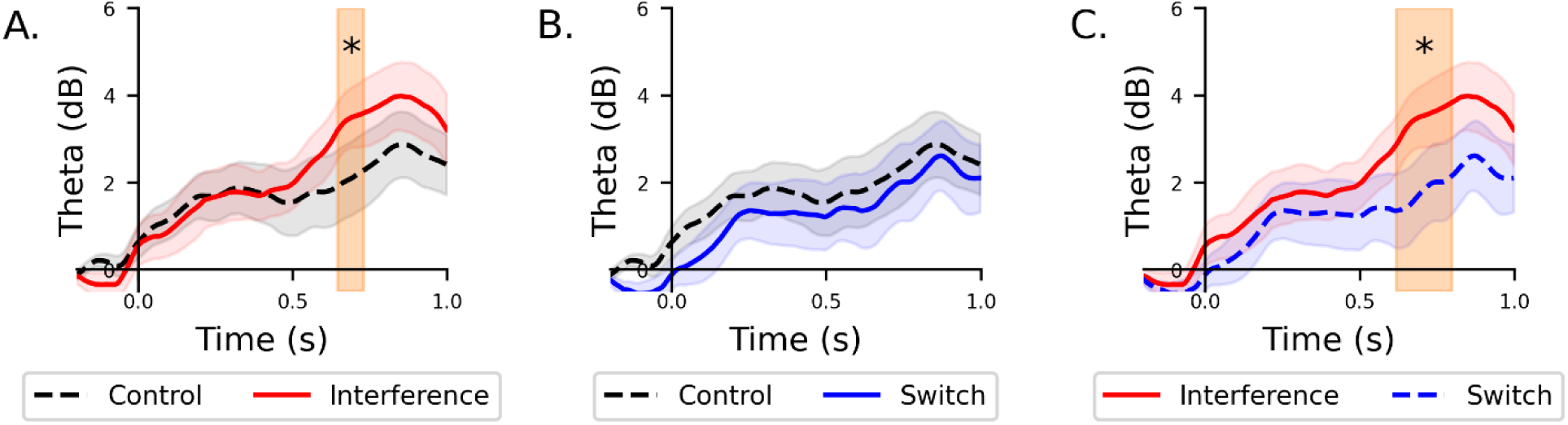
Theta-band EEG power in the frontal region. Change in theta-band EEG power relative to the stimulus onset, expressed in dB, for interference and control conditions (A), switch and control conditions (B), and interference and switch conditions (C). Theta power was computed within the 3.5–7.5 Hz frequency range and averaged across frontal electrodes (F3, Fz, F4). Significant differences were identified using a cluster-based permutation test, and the orange shaded regions highlight significant time intervals. Notably, theta power was significantly enhanced in the interference condition compared to both the control and switch condition. Also, there is no significant difference between control and switch conditions.

## Discussion

Typically, task-switching and set-shifting paradigms involve incongruent bivalent stimuli and report both shifting and interference control as aspects of cognitive flexibility (e.g, Wylie & Allport, 2000; Monsell et al., 2000). Our research aims to determine how cognitive flexibility without interference control in response level is manifested in the brain’s activity. In particular, we seek to identify distinctive EEG correlates of shifting and interference control, thereby providing insights into the underlying brain processes. Both conditions were tested in a single experiment, using consistent materials, to ensure that the remaining differences directly reflect the distinct cognitive processes involved in cognitive flexibility and interference control.

Behavioral analysis showed a delay in RT of approximately 55 ms in task switch condition compared to control, consistent with the literature and referred to as switch cost (Jersild, 1927). Our results further highlight the importance of the frontal region in cognitive flexibility tasks. We observed N2 in the centro-frontal region during the switch condition and decreased amplitude in P3a in the frontal region. This finding aligns with the established role of the frontal site in tasks requiring cognitive flexibility (Milner, 1963; Monsell, 2003; Aron et al., 2004). Also, consistent with evidence indicating that patients with left frontal lobe damage exhibit behavioral abnormalities when switching between attributes (Keele & Rafal, 2000; Rogers et al., 1998; Monsell, 2003).

Previous research has frequently reported increased fronto-central N2 amplitude and reduced parietal P3b in switch trials compared to repeat trials (Kopp et al., 2020; Grundy et al., 2013; Lavric et al., 2008; Gajewski & Falkenstein, 2011; Karayanidis et al., 2003). However, our study just found significant N2 modulation. This may be due to the use of univalent stimuli, which are associated with reduced task difficulty and smaller switch costs (Elchlepp et al., 2012). As P3b amplitude is linked to task difficulty (e.g., Isreal et al., 1980), the relative ease of our task likely diminished the P3b effect. Further, studies using bivalent stimuli have shown larger valence effects on P3 during switch trials (Hsieh & Liu, 2008; Karayanidis et al., 2003), suggesting P3 might be more related to stimulus valence than task switching itself (Karayanidis & Jamadar, 2014). The absence of a P3b effect in our findings, therefore, likely reflects the distinct cognitive processes indexed by N2 and P3b, combined with the lower task demands associated with univalent stimuli. This contrasts with previous research that often employed incongruent bivalent stimuli (e.g, Wylie & Allport, 2000; Monsell et al., 2000), potentially confounding interference control and cognitive flexibility demands.

In terms of interference control, behavioral analysis showed a delay in RT of approximately 167 ms in the interference condition compared to the control, consistent with the literature and referred to as interference effect (Stroop, 1935). In ERP analysis, we observed increased N400 amplitude in the parietal area and a positive deviation in the occipital region during the interference condition compared to the control in ERP analysis. This aligns with previous research (Donohue et al., 2016; Hanslmayr et al., 2008; Larson et al., 2009; Liotti et al., 2000; Markela-Lerenc et al., 2004; Qiu et al., 2006; Rebai et al., 1997). The posterior N400 has also been identified in tasks demanding conflict management, indicating diverse processes such as conflict monitoring and control execution, specifically interference suppression (Heidlmayr et al., 2020). Additionally, theta band activity was enhanced in the frontal region (over mid-frontal electrodes) after stimulus presentation, consistent with previous evidence supporting its role in interference control during the Stroop task, particularly in conflict processing and conflict detection (Hanslmayr et al., 2008; Ergen et al., 2014; Cohen, 2014; Ridderinkhof et al., 2004; Eschmann & Mecklinger, 2021; Haciahmet et al., 2023).

Direct comparison between interference and switch conditions revealed that the N2 component in the centro-frontal region was significantly larger in the switch condition rather than the interference condition, while the theta activity in the frontal region was prominent in the interference condition. Based on prior studies, both N2 in the frontal site and theta oscillation in mid frontal site are linked to conflict monitoring (e.g. Cohen, 2014; Heidlmayr et al., 2020). The increased midfrontal theta activity observed during the interference condition and the pronounced N2 component in the switch condition indicate that distinct neural mechanisms are involved in conflict monitoring for within-task interference and task switching. The increased theta in interference condition likely reflects a sustained demand for cognitive control to manage interference and maintain task performance. Conversely, the N2 in switch trials points to phasic conflict detection and inhibitory processes at the task-set level, possibly reflecting the suppression of the previous task set or the resolution of competing task rules. Moreover, prior studies have attributed theta activity and the N2 component in the mid-frontal region to the anterior cingulate cortex (Cohen, 2014; Heidlmayr et al., 2020). Based on our data, it appears that this region plays a critical role in executive functions such as cognitive flexibility and interference control, though it serves distinct functions within each process.

Additionally, we observed a positive LSP in the occipital region for both switch and interference conditions. This finding is consistent with prior research suggesting that the LSP reflects engagement of central executive processes (Hanslmayr et al., 2008). Furthermore, it supports the “unity and diversity” model of executive functions, which posits that these functions are composed of distinct components with unique neural signatures while also being interconnected and sharing overlapping processes (Teuber, 1972; Friedman et al., 2008; Miyake & Friedman, 2013). The LSP occurred earlier in the switch condition than in the interference condition, mirroring differences in reaction times, which were also faster in the switch condition. This temporal difference suggests that the LSP may be related to response mechanisms, potentially reflecting processes linked to response preparation or execution.

## Acknowledgements

This research was funded by the InnoBrain AB company and partially funded by the Swedish Research Council through grant agreement no. 2018-05973, KTH Royal Institute of Technology and Digital Futures.

